# Dwarf open reading frame (DWORF) peptide is a direct activator of the sarcoplasmic reticulum calcium pump SERCA

**DOI:** 10.1101/2020.10.01.322610

**Authors:** M’Lynn E. Fisher, Elisa Bovo, Ellen E. Cho, Marsha P. Pribadi, Michael P. Dalton, M. Joanne Lemieux, Nishadh Rathod, Rodrigo Aguayo-Ortiz, L. Michel Espinoza-Fonseca, Seth L. Robia, Aleksey V. Zima, Howard S. Young

## Abstract

The cardiac sarcoplasmic reticulum calcium pump, SERCA, sequesters calcium in the sarco-endoplasmic reticulum (SR/ER) and plays a critical role in the contraction-relaxation cycle of the heart. A well-known regulator of SERCA in cardiac muscle is phospholamban (PLN), which interacts with the pump and reduces its apparent calcium affinity. A newly discovered SERCA regulatory subunit in cardiac muscle, dwarf open reading frame (DWORF), has added a new level of SERCA regulation. In this report, we modeled the structure of DWORF and evaluated it using molecular dynamics simulations. DWORF structure was modeled as a discontinuous helix with an unwound region at Pro^15^. This model orients an N-terminal amphipathic helix along the membrane surface and leaves a relatively short C-terminal transmembrane helix. We determined the functional regulation of SERCA by DWORF using a membrane reconstitution system. Surprisingly, we observed that DWORF directly activated SERCA by increasing its turnover rate. Furthermore, in-cell imaging of calcium dynamics demonstrated that DWORF increased SERCA-dependent ER calcium load, calcium reuptake rate, and spontaneous calcium release. Together, these functional assays suggest opposing effects of DWORF and PLN on SERCA function. The results agree with fluorescence resonance energy transfer experiments, which revealed changes in the affinity of DWORF for SERCA at low versus high cytosolic calcium concentrations. We found that DWORF has a higher affinity for SERCA in the presence of calcium, while PLN had the opposite behavior, a higher affinity for SERCA in low calcium. We propose a new mechanism for DWORF regulation of cardiac calcium handling in which DWORF directly enhances SERCA turnover by stabilizing the conformations of SERCA that predominate during elevated cytosolic calcium.

## INTRODUCTION

The sarco-endoplasmic reticulum calcium pump (SERCA) is an ion transporting ATPase that plays a critical role in intracellular calcium signaling. SERCA maintains calcium content of the sarco-endoplasmic reticulum, creating a 2000-fold gradient that can be mobilized for signaling via inositol-1,4,5-triphosphate receptor (IP3R) and ryanodine receptor (RyR) calcium channels. This signaling system is essential for normal cell physiology, and disordered calcium handling underlies a diverse array of diseases including cardiomyopathies (1), skeletal muscle disorders (2), and neurological diseases (3). In the heart, the cardiac-specific isoform SERCA2a maintains sarcoplasmic reticulum (SR) calcium, which is the major source of calcium for cardiac contractility. While SERCA calcium transport is essential for all cells, it is particularly important in cardiomyocytes where defects in SERCA activity or regulation are associated with cardiomyopathies and heart failure (4–6). This connection with heart failure has focused attention on SERCA as a possible target for therapy. An initially promising approach in animal models, increasing SERCA expression via gene delivery (7), has proven challenging in human clinical trials (8). Nonetheless, the rationale for treating heart failure by improving calcium transport function remains compelling and orthogonal approaches may be required.

The regulation of SERCA in normal cellular calcium homeostasis is of fundamental importance, and dysregulation is a major mechanism for disease development and progression. In the heart, SERCA activity is regulated to allow dynamic calcium homeostasis, which changes in response to the need for cardiac output during rest or exertion. A small transmembrane protein, phospholamban (PLN), is the primary regulatory subunit of SERCA in cardiac muscle. PLN inhibits SERCA by decreasing its apparent calcium affinity, thereby reducing both the rate of calcium reuptake and the amount of calcium in the SR. PLN inhibition of SERCA controls the rate of relaxation and the SR calcium load for subsequent contractions, modulating both the lusitropic and inotropic properties of the heart. PLN inhibition of SERCA is relieved by phosphorylation with the main mechanism involving protein kinase A (PKA) and the β-adrenergic pathway (9). Under resting conditions, PLN is an important brake that prevents SR calcium overload and the arrhythmogenic consequences (10). In turn, regulation of PLN by phosphorylation – regulation of the regulator – creates a dynamic SR calcium load that can respond to the sympathetic need for increased cardiac output, the so-called “fight-or-flight” response. Thus, there exists a pool of SERCA pumps in the SR membrane and their combined calcium transport capacity can be finely tuned – upregulated or downregulated – depending on the requirement for cardiac contractility.

Since the initial discovery of PLN decades ago (9), it has stood as the only regulatory subunit of SERCA in ventricular muscle. This changed recently with the discovery of a small transmembrane SERCA-binding protein, dwarf open reading frame (DWORF) (11). DWORF has weak sequence similarity to PLN and its skeletal muscle homolog sarcolipin (SLN), and it appears to comprise a transmembrane peptide of unknown structure and function (**Figure 1**). DWORF was found to increase SERCA activity by opposing PLN function, leading to the hypothesis that DWORF is a non-inhibitory competitor that displaces PLN from the inhibitory groove of SERCA (12). This raised the question, why is an additional means of reversing PLN inhibition necessary? PLN inhibition can be reversed by phosphorylation via PKA, CaMKII (13), and Akt (14), as well as by elevated calcium concentrations. With this level of apparent redundancy in reversing SERCA inhibition by PLN, which now presumably includes DWORF, a detailed study of the relative control of SERCA by DWORF was required. Toward this goal, we modeled the structure of DWORF and evaluated its properties using molecular dynamics simulations. We then compared the ability of DWORF and PLN to directly regulate SERCA function. To determine the effects of DWORF on SERCA in vivo, we measured SERCA-dependent calcium dynamics and SERCA binding by these regulatory peptides during active intracellular calcium signaling. The results provide insight into DWORF structure and function and a role for DWORF in the direct regulation of the SERCA calcium pump.

**Figure 1.**
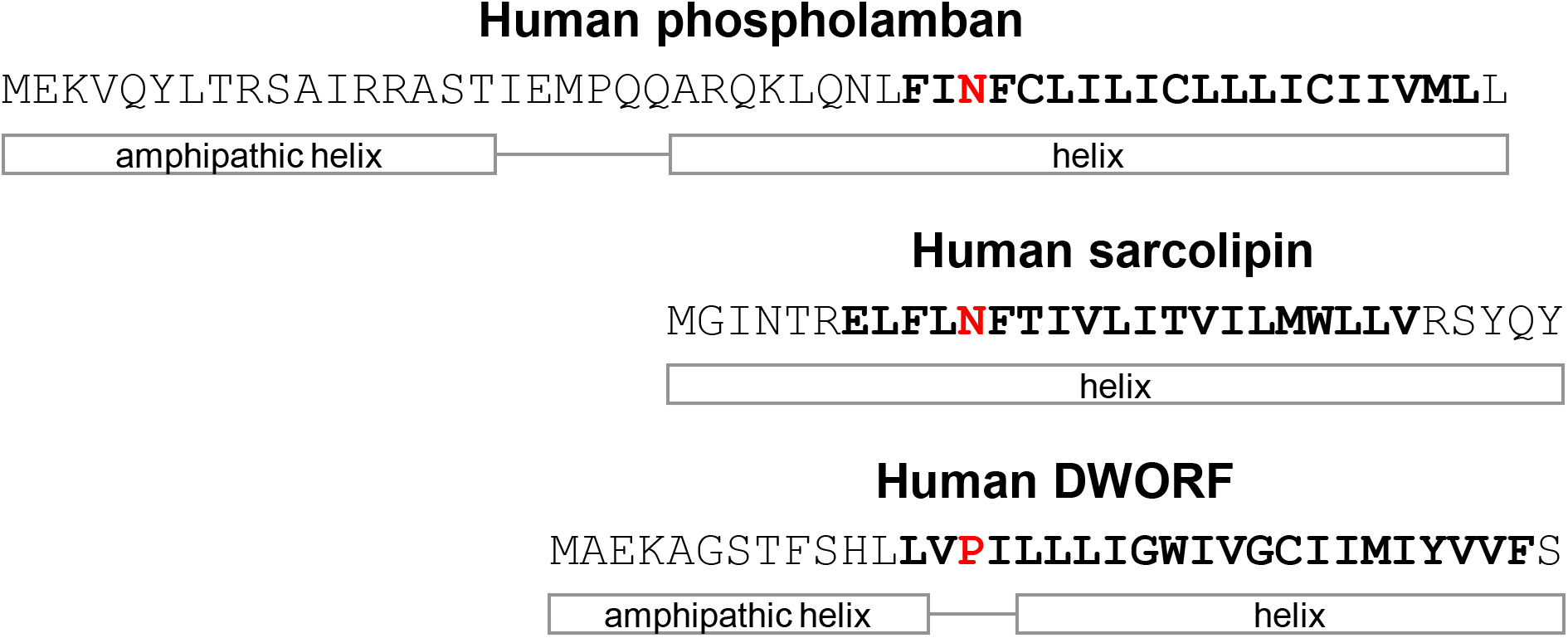
**Figure legend:** Sequence alignments, secondary structure predictions, and transmembrane domains for phospholamban, sarcolipin, and DWORF. The predicted transmembrane domains are in bold letters. The sequences are aligned between Asn^34^ of phospholamban, Asn^11^ of sarcolipin, and Pro^15^ of DWORF. The helical regions of phospholamban are based on the NMR structure (PDB code 2KYV). The helical region of sarcolipin is based on the X-ray crystal structures of the SERCA complex (PDB codes 4H1W & 3W5B). The helical regions of DWORF are based on sequence predictions and molecular dynamics simulations in the present study.

## RESULTS

### Structure of DWORF

In the present work, we used molecular modeling to generate two structures of DWORF, one modeled as a continuous α-helix and another modeled as an N-terminal α-helix (residues 1-13), a flexible linker (residues 14-16, centered around Pro^15^), and a C-terminal α-helix (residues 17-35). The models were embedded in a lipid bilayer and equilibrated using molecular dynamics (MD) simulations for 2000 ns (**Figure 2A**). Both starting models quickly converged on a helix-linker-helix structure, though the overall structure was dynamic in the simulations and varied somewhat depending on the starting model. A similar behavior has been observed with SLN (15). We consider the second model, the helix-linker-helix model with an unwound region centered on Pro^15^, to be the more stable and probable structure of DWORF (**Figure 2B**). This model was maintained over 2000 ns of MD simulations, though again the structure was dynamic on its own in a lipid bilayer (**Figure 2A**). To further rationalize this model, we carried out predictions of the secondary structure (16–19) and the transmembrane region (16,20–22) of the DWORF peptide using only the amino acid sequence (**Figure 1**). The secondary structure prediction suggested that the N-terminal region (residues 1-8) has a high probability of being found as a random coil, while the remainder of the peptide (residues 9-35) is likely to be found as an α-helix. Prediction of the transmembrane domain showed that this region comprises approximately residues 10 to 33, which coincided with the predicted α-helical region. These predictions appeared inconsistent with either starting model.

**Figure 2.**
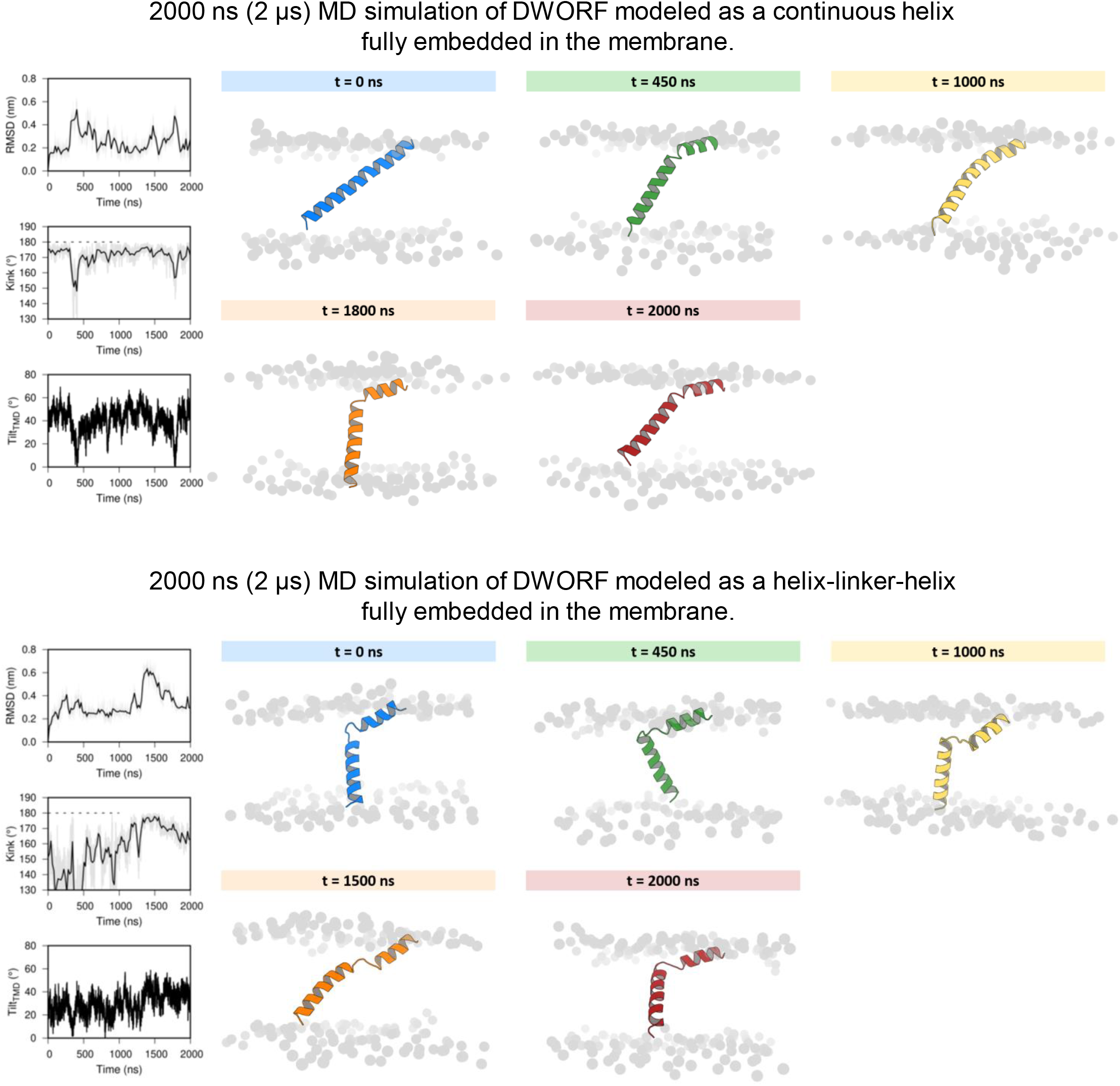
**Figure legend:** Helix-linker-helix molecular model of DWORF. (**A**) 2000 ns MD simulations of DWORF modeled as a continuous helix (top panels) and a helix-linker-helix (bottom panels). Shown are snapshots during the simulations as well as the RMSD, kink angle, and tilt angle of the transmembrane helix TM. Notice that both simulations support a helix-linker-helix structure of DWORF. The MD simulation in the lower panel indicates that the helix-linker-helix structure is maintained, though it is dynamic during the time course of the simulation.

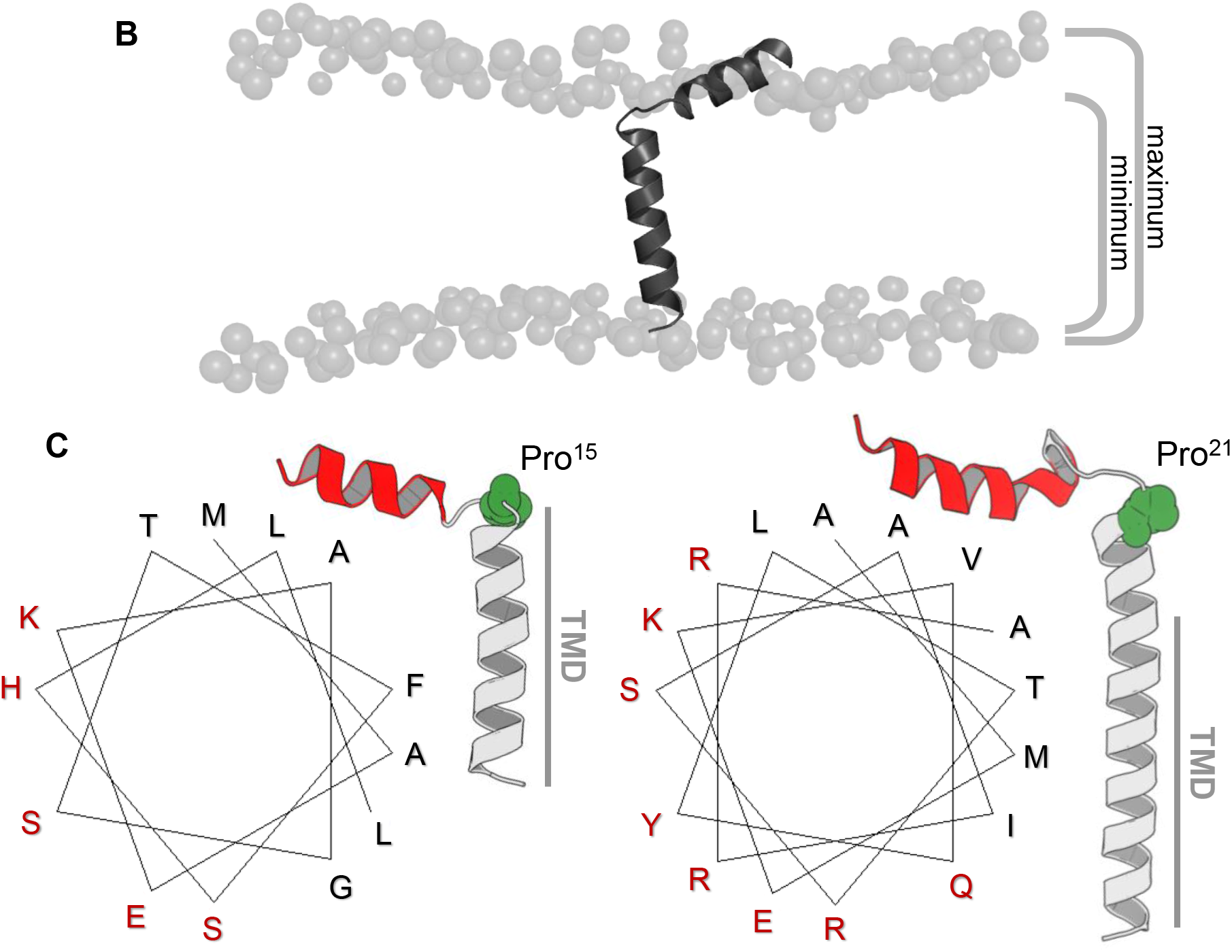
**Figure legend:** Helix-linker-helix molecular model of DWORF. (**B**) Molecular dynamics simulation of DWORF in a lipid bilayer. DWORF was modeled as an N-terminal helix, a kink centered around Pro^15^, and a C-terminal transmembrane helix. The phospholipid headgroups are indicated as spheres. Notice how the bilayer becomes thinner around the short transmembrane helix of DWORF. The maximum and minimum widths of the lipid bilayer are indicated. (**C**) Hydrophobic moment analysis reveals amphipathic helices at the C-terminus of DWORF and PLN. Notice the similar distribution of polar (red) and apolar (black) residues in the helix-kink structure of DWORF and PLN N-terminal to a proline residue (Pro^15^ in DWORF and Pro^21^ in PLN).

Nonetheless, the existence of the N-terminal juxtamembrane helix in the second model was preferred because of the “helix breaker” property of the proline residue located at position 15. Therefore, we carried out a hydrophobic moment analysis to understand the structural behavior of this N-terminal region (**Figure 2C**). Hydrophobic moment analysis revealed an amphipathic helix comprising is remarkably like the N-terminus of PLN, though DWORF has a shorter sequence following the proline residue (Pro^21^ in PLN), which includes only the transmembrane domain. In PLN, a polar helical region (residues Gln^22^ to Asn^30^) follows the proline residue and precedes the transmembrane domain (residues 31-51) and this feature is absent in DWORF. High hydrophobic moment values indicated that the presence of this helix allows hydrophobic residues to be located on one side of the helix and hydrophilic residues on the other. In the molecular model, the hydrophobic residues are oriented towards the lipid bilayer, while the hydrophilic residues are exposed to the aqueous solvent. This is also observed in the juxtamembrane regions of PLN (PDB code 2KYV (23)) and the amyloid precursor protein (PDB code 2LLM (24)). Despite the result of the secondary structure prediction, the formation of this juxtamembrane helix suggests that DWORF may mimic some of the structural features of PLN.

### SERCA activity in the presence of DWORF

The unique structure and puzzling role proposed for DWORF (competitive inhibitor of an inhibitor (11)), prompted us to look for a direct functional effect of DWORF on SERCA. To achieve this, DWORF was co-reconstituted into proteoliposomes with SERCA and the calcium-dependent ATPase activity was measured (**Figure 3**). For comparison, the previously reported ATPase activity curves for SERCA in the absence and presence of PLN and SLN are shown in **Figure 3A** (25,26) and the data for SERCA in the absence and presence of DWORF are shown in **Figure 3B**. At a molar ratio of ~1-2 DWORF per SERCA, DWORF increased the turnover rate of SERCA at nearly all calcium concentrations tested (0.01 to 15 μM free calcium). At this DWORF-SERCA ratio, the maximal activity (V_max_) of SERCA in the absence of DWORF increased 1.7-fold in the presence of DWORF (V_max_ values were 4.1 ± 0. and 6.9 ± 0.1 μmoles/min/mg, respectively). This level of SERCA activation is comparable to the small molecule activator CDN1163 (27). There was no significant effect of DWORF on the apparent calcium affinity of SERCA (K_Ca_ values of 0.42 ± 0.03 and 0.48 ± 0.03 μM calcium, respectively). At higher concentrations of DWORF, ~5-7 DWORF per SERCA, DWORF acted as an inhibitor of SERCA with characteristics similar to SLN (**Table 1**). The V_max_ of SERCA decreased to 3.0 ± 0.1 μmoles/min/mg in the presence of excess DWORF and there was an inhibitory effect of DWORF on the apparent calcium affinity of SERCA (K_Ca_ value increased from 0.42 ± 0.03 to 0.81 ± 0.04 μM calcium in the presence of DWORF). Thus, equimolar levels of DWORF caused a direct activation of SERCA, which shifted to a SLN-like inhibitory effect with excess DWORF. We consider the equimolar condition to be physiological and the excess condition to be non-physiological (11).

**Figure 3.**
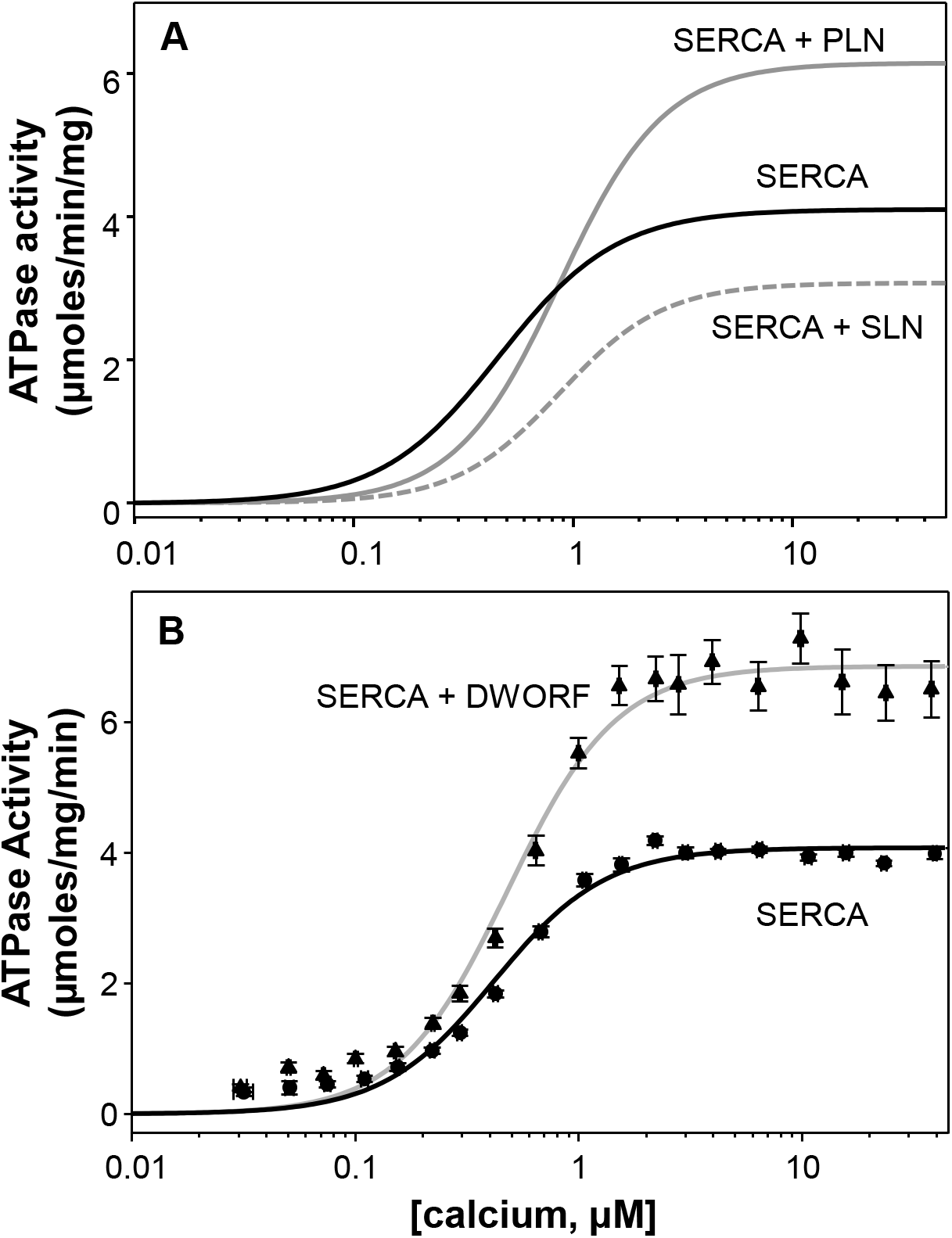
**Figure legend:** ATPase activity of SERCA co-reconstituted in the absence and presence of DWORF. **(A)** Representation of ATPase activity of SERCA in the absence (black line) and presence of phospholamban (gray line) and sarcolipin (gray dashed line). **(B)** Topology diagrams for phospholamban, sarcolipin, and DWORF. The cytoplasmic domains are in white, the transmembrane domains in gray, and the luminal domains in black. An asparagine residue critical for PLB and SLN function is indicated, as is the corresponding proline residue in DWORF.

### Cellular SERCA calcium transport in the presence of DWORF

Co-reconstitution of SERCA with DWORF resulted in increased ATPase activity (**Figure 3**). Compared to previous reports (12), we observed a direct activation of SERCA by DWORF, even in the absence of PLN. The data suggested that DWORF not only relieves SERCA inhibition by displacing PLN, but it directly stimulates SERCA activity. To determine whether increased ATPase activity was accompanied by increased cellular calcium transport activity, we utilized a newly developed approach for measuring calcium uptake into the endoplasmic reticulum of live cells (28). The technique used an inducible human SERCA2a stable cell line (t-Rex-293 cells) with the calcium release channel RyR2 (allows manipulation of ER calcium load), and an ER-targeted calcium indicator R-CEPIA1er (allows measurement of ER calcium load) (28,29). The cell line was transiently transfected with mCer-DWORF (or mCer-PLN) to assess regulation of SERCA2a. mCer-DWORF was predominantly expressed in the ER (estimated from the overlap between the mCer-DWORF and the R-CEPIA1er signals; **Figure 4A**), with a distribution pattern similar to SERCA (28). To determine SERCA2a function, the plasma membrane was selectively permeabilized with saponin to allow control of the cytosolic environment, including free calcium and ATP concentrations. ER calcium uptake was quantified from the increase in R-CEPIA1er fluorescence after full ER calcium depletion by caffeine followed by RyR2 inhibition with ruthenium red and tetracaine (RyR+Tetr; **Figure 4B**), which blocks the principal calcium leak pathway (28). The first derivative of ER calcium uptake (*d*[Ca^2+^]_ER_/*d*t) was plotted against the corresponding [Ca^2+^]_ER_ to estimate the maximum ER calcium uptake rate and the maximum ER calcium load (**Figure 4C**). As expected, PLN transfection decreased ER calcium uptake compared to control. The opposite effect was observed with DWORF transfection, which almost doubled SERCA calcium uptake rate over the entire range of physiological ER calcium loads. These data indicated that DWORF can act as a potent activator of SERCA2a and this regulation is direct, not requiring pre-existing inhibition of SERCA by PLN. Moreover, it was noteworthy that DWORF enhanced the maximum ER calcium load (**Figure 4C, arrow**). Since the thermodynamic driving force for calcium transport is the ATP/ADP ratio, we conclude that DWORF enhances SERCA catalytic efficiency (energetic cost of calcium transport) (28).

**Figure 4.**
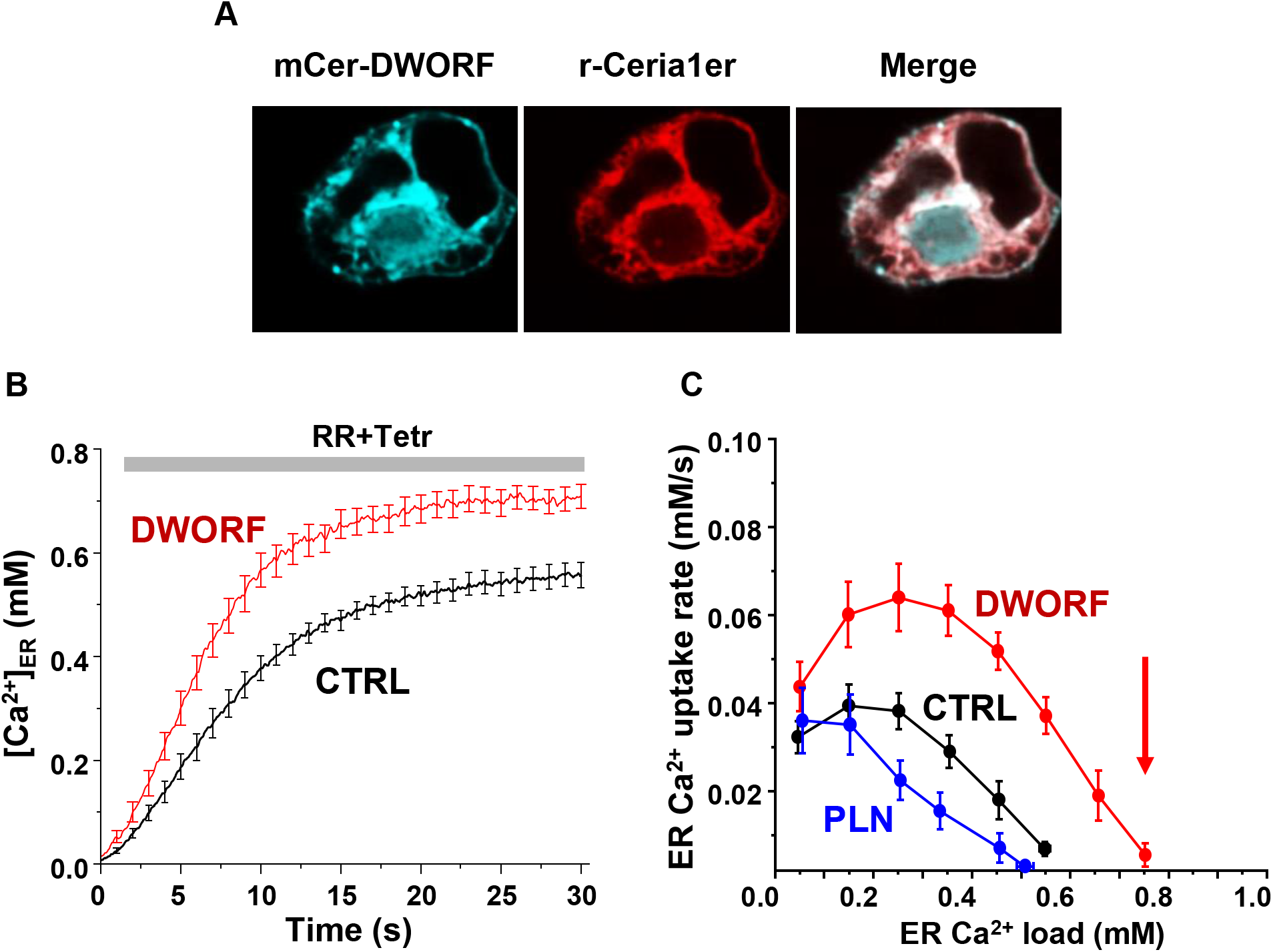
**Figure legend:** DWORF enhances SERCA-dependent calcium dynamics *in vivo*. **(A)** Inducible human SERCA2a stable cell line (t-Rex-293 cells + Tetr) were transfected with mCer-DWORF together with the Ca^2+^ release channel RyR2 and the ER-targeted Ca^2+^ indicator R-CEPIA1er. MCer-DWORF expression in these cells showed a similar pattern as R-CEPIA1er. **(B)** The rate of [Ca^2+^]ER reuptake after full ER Ca^2+^ depletion by caffeine followed by RyR2 inhibition with ruthenium red (RR). **(C)** The ER Ca^2+^ uptake rate was plotted against the corresponding ER [Ca^2+^] load. DWORF expression almost doubled ER Ca^2+^ uptake rate through the entire range of physiological ER Ca^2+^ loads, which is the opposite trend seen for the SERCA inhibitor PLN. DWORF also increased the ER [Ca^2+^] load (arrow).

To investigate the effect of DWORF on SERCA in cellular calcium handling dynamics, we mimicked cardiac calcium handling in a heterologous cell model expressing SERCA and RyR (**Figure 5**). This system generates periodic calcium waves due to spontaneous calcium release by RyR2 followed by calcium reuptake by SERCA (**Figure 5A,B**). Co-expression of DWORF resulted in an increased frequency and amplitude of spontaneous calcium waves, suggesting a significantly increased ER calcium load and faster calcium reuptake during calcium waves due to increased SERCA activity. In contrast, PLN significantly decreased the calcium uptake rate by slowing the calcium wave recovery (**Figure 5B**). Similar effects of DWORF on spontaneous calcium release were observed when calcium waves were measured as cytosolic calcium fluctuation in intact cells (**Figure 5C**) with a significant increase in the average amplitude and integral of RyR-mediated calcium release events in DWORF expressing cells (**Figure 5D**). These results demonstrate that DWORF expression increases ER calcium load by facilitating SERCA mediated calcium uptake. These data are consistent with the unique structure of DWORF (**Figure 2**) and the observation that DWORF directly increases the calcium-dependent ATPase activity of SERCA (**Figure 3**).

**Figure 5.**
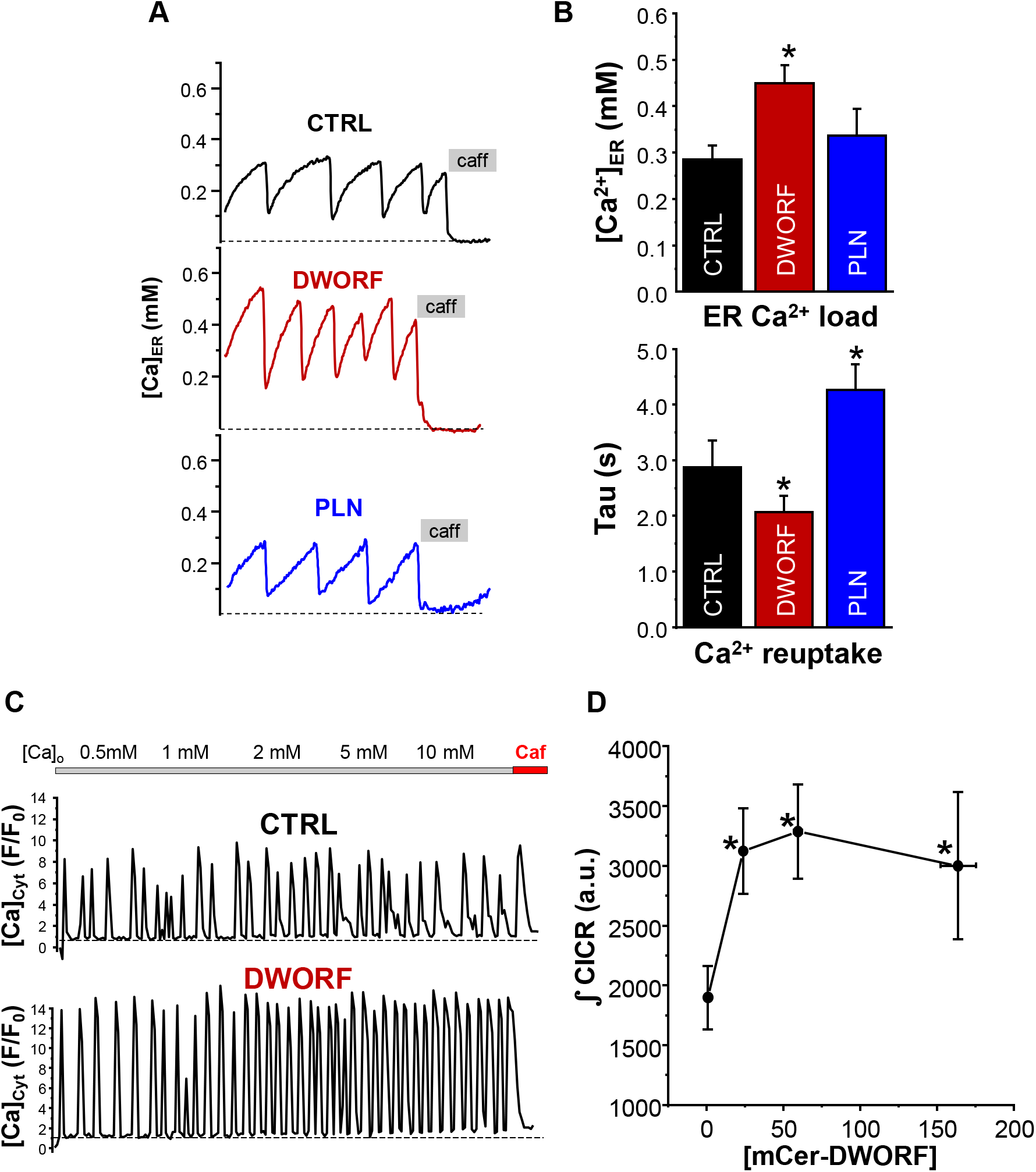
**Figure legend:** Effect of DWORF on calcium induced calcium release (CICR). **(A)** and **(B)** Co-expression of SERCA2a and RyR2 produced Ca^2+^ waves due to spontaneous activation of RyR2 followed by SERCA Ca^2+^ reuptake. DWORF had the tendency to increase the magnitude and frequency of spontaneous Ca^2+^ waves, while PLN significantly decreased it. DWORF had the tendency to increase the occurrence of spontaneous Ca^2+^ waves, while PLN significantly decreased it. Similar effects of DWORF on CICR were observed when Ca^2+^ waves were measured as cytosolic Ca^2+^ fluctuation in intact cells. (**C**) Average amplitude and frequency of RyR-mediated Ca^2+^ release events were significantly increased in DWORF expressing cells. (**D**) The integral of RyR-mediated Ca^2+^ release events was significantly increased in DWORF expressing cells.

### Dynamics of DWORF binding to SERCA

To gain insight into the mechanistic differences in the regulation of SERCA by PLN and DWORF, we measured the binding of these regulins to SERCA in a cell membrane. Fluorescent protein tags were fused to SERCA, PLN, and DWORF and intermolecular FRET was quantified in cells expressing pairs of fusion proteins. This method was recently used to show that PLN and DWORF had similar apparent affinities for SERCA in intact cells (12,30). However, we previously showed that PLN affinity for SERCA is sensitive to calcium, where the affinity was decreased by micromolar calcium in permeabilized cells or during extended calcium elevations in rapidly paced cardiac myocytes (31). This apparent change in affinity suggests that PLN binds more avidly to the conformations of SERCA that prevail under resting conditions when cytosolic calcium is low. The enhanced interaction with calcium-free forms of SERCA offers a mechanistic explanation for the effect of PLN on the apparent calcium affinity of SERCA. PLN binds to SERCA in a groove formed by transmembrane segments M2, M6, and M9. Upon calcium binding, M2 undergoes a large conformational change that closes the inhibitory groove and forms the calcium bound E1 state of SERCA. PLN appears to act as a competitive inhibitor of calcium binding by impeding groove closure and the E2-E1 transition (32–34).

To determine how the SERCA-DWORF regulatory complex responds to changes in calcium, we permeabilized cells expressing Cer-SERCA and YFP-PLN or YFP-DWORF (**Figure 6A, B**) in solutions mimicking low (diastolic; 100 nM) or high (systolic; 3 μM) intracellular calcium concentrations. FRET was quantified for each cell and compared to that cell’s level of expression of the YFP acceptor. As previously observed (35), FRET from SERCA-PLN was lowest for cells expressing low levels of protein and increased to a maximal level of ~25% FRET efficiency for high expressing cells (**Figure 6A**). The relationship was well-described by a hyperbolic fit that yielded the apparent dissociation constant (K_D_) of the SERCA-PLN complex (**Figure 6C**). Permeabilization of cells in high calcium yielded a binding curve that was right-shifted to higher protein concentrations, suggesting a decrease in SERCA-PLN binding affinity compared to low calcium conditions (31). In contrast, the SERCA-DWORF regulatory complex showed the opposite response to calcium, with a small left-shift of the hyperbolic FRET versus protein concentration curve (**Figure 6B**). Replicate experiments are summarized in **Figure 6C**. The data indicate that high calcium decreases the affinity of PLN for SERCA, yet it has the opposite effect on DWORF where it increases the affinity for SERCA. These data are in agreement with functional measurements that show DWORF directly increases SERCA activity (**Figure 3**), as well as SERCA transport kinetics and thermodynamics in cells (**Figures 4** & **5**).

**Figure 6.**
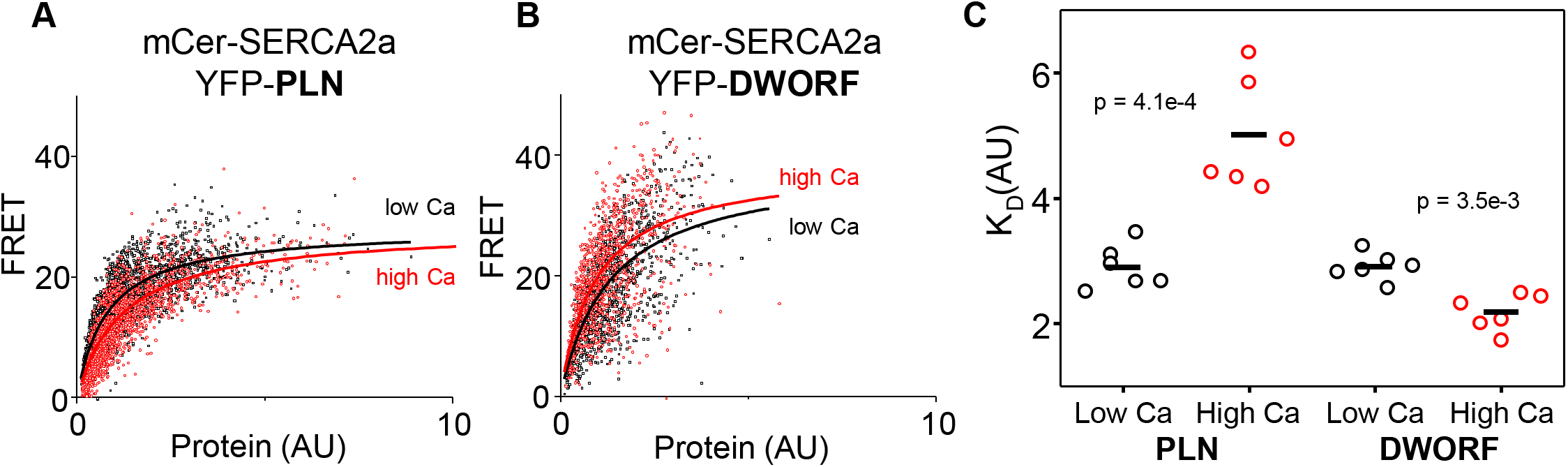
**Figure legend:** FRET analysis of SERCA-DWORF interactions. The average acceptor sensitization FRET efficiency of cells co-transfected with mCer-SERCA2a and either **(A)** YFP-PLN or **(B)** YFP-DWORF. FRET efficiency was measured at high and low calcium concentrations to assess the relative affinity of PLN and DWORF for the calcium-free and calcium-bound conformations of SERCA. **(C)** Hyperbolic fits to data provide quantification of the apparent dissociation constant (K_D_) of the SERCA-PLN or SERCA-DWORF regulatory complexes. Ca decreases the apparent affinity of PLN for SERCA, yet it increases the affinity of DWORF for SERCA.

## DISCUSSION

### Functional effect of DWORF

The data presented here demonstrate that DWORF has a direct functional effect on SERCA, which enhances calcium-dependent ATPase activity of SERCA in an isolated system and enhances SERCA-dependent calcium dynamics in living cells. While DWORF may displace PLN from SERCA as previously suggested (11), we have shown that DWORF also has a direct effect on SERCA function that is opposite to the inhibitory properties of PLN. The opposing functions of DWORF and PLN represent a previously unknown regulatory axis for fine-tuning SERCA-dependent calcium homeostasis in response to demand for cardiac output.

The reconstitution-based system allowed us to assess the effect of DWORF on SERCA function in an isolated, controlled membrane environment. This SERCA-DWORF two-component system is well-defined and finely controlled to allow for measurement of ATPase activity (26,36–39) or calcium transport (40) in the presence of regulatory subunits such as PLN, SLN, and DWORF. PLN and SLN are known inhibitors of SERCA in that they reduce its apparent affinity for calcium (**Figure 3A**). PLN has also been shown to increase the maximal activity of SERCA, while SLN has been shown to decrease the maximal activity. Herein, we measured SERCA ATPase activity in the presence of DWORF. At a near equimolar ratio, DWORF acted as a direct activator by increasing the maximal activity of SERCA 1.7-fold without an effect on the apparent calcium affinity (**Figure 3B**). It is interesting that the two known regulators of SERCA in cardiac muscle, DWORF and PLN, both increase the maximal activity of SERCA. The magnitude of the V_max_ increase seen in the presence of DWORF (~1.7-fold) is similar to that seen in the presence of excess PLN (~1.5-fold (25)) and a small molecule activator CDN1163 (~1.5-fold (27)). In contrast, SLN in skeletal muscle decreases the maximal activity of SERCA. The important features of this DWORF-PLN dual-peptide regulation in cardiac muscle is that the regulatory subunits have distinct functions. The primary effect of PLN is to alter the calcium affinity of SERCA and the primary effect of DWORF is to alter the maximal activity of SERCA. PLN can also increase the maximal activity of SERCA, but this occurs only under conditions of excess PLN to SERCA (25). If correct, these different but complimentary functions would allow for a fine level of rheostatic control of SERCA-mediated calcium homeostasis.

### Cellular effect of DWORF

Membrane co-reconstitution systems can provide mechanistic insight into SERCA regulation; however, it is both an advantage and disadvantage that they lack the complexity of a cellular environment. HEK293 cells have been established as a novel cardiomimetic system for evaluating SERCA-dependent calcium-handling and its regulation by PLN (28). This approach avoids the confounding effects of imaging cells during contractions. Co-expression of SERCA2a and RyR2 in this cell system causes periodic calcium transients (waves) similar to those that elicit contractions in cardiomyocytes. The HEK cell model lacks endogenous SERCA regulators and it can be stably transfected with exogenous regulators such as PLN and DWORF. This enabled the observation of ER calcium load, calcium uptake rate, and changes in the magnitude and frequency of calcium waves. These are SERCA-dependent cellular responses to changes in calcium concentration and the presence of SERCA regulatory subunits. This is a well-defined model system for testing the effects of regulatory peptides such as DWORF on SERCA-dependent cellular calcium dynamics. As was observed for PLN, DWORF co-localized with SERCA in ER membranes of HEK293 cells (**Figure 4A**). DWORF increased two key parameters of SERCA function, ER calcium content (**Figure 3B**) and ER calcium uptake rate (**Figure 3C**). These effects were opposite to what is seen for PLN. The advantage of this approach is that it enabled the examination of SERCA function in a cellular system in terms of the maximum calcium uptake rate and maximum calcium load of the ER. These parameters reflect the kinetics and thermodynamics of SERCA in the absence and presence of peptides such as DWORF.

An alternative measure of ER calcium load and SERCA activity is provided by a cardiomimetic HEK system that generates periodic calcium waves due to spontaneous calcium release by RyR2, followed by SERCA calcium reuptake. Calcium release via RyR2 is known to depend on ER calcium load, which is determined by SERCA activity. Thus, the cytosolic calcium concentration during periodic calcium waves is an indirect measure of SERCA function in the absence and presence of SERCA regulatory peptides (**Figure 5A, B**). DWORF increased the amplitude and frequency of RyR-mediated calcium release events (**Figure 5C, D**). The SERCA-mediated calcium reuptake rate, ER calcium load, and calcium waves were all increased in DWORF expressing cells, strongly suggesting that DWORF directly activates SERCA and increases its catalytic efficiency. These cellular data (**Figures 4 & 5**) are in agreement with the activity data from reconstituted proteoliposomes containing SERCA and DWORF (**Figure 3**).

This raised the question, what is the mechanism by which DWORF activates SERCA? To address this, FRET efficiency between SERCA-PLN and SERCA-DWORF was measured at high and low calcium concentrations to assess the relative affinity of PLN and DWORF for the calcium-bound and calcium-free conformations of SERCA (**Figure 6A, B**). The data provided relative measures of the dissociation constants (K_D_) of the SERCA-PLN and SERCA-DWORF regulatory complexes. As previously observed, elevated calcium did not abolish binding of PLN to SERCA, instead it reduced the apparent affinity of PLN for SERCA. In contrast, calcium increased the apparent affinity of DWORF for SERCA (**Figure 6C**), revealing that DWORF prefers to interact with conformations of SERCA that prevail at high calcium. In the context of the current paradigm of DWORF as a competitive inhibitor of PLN binding to SERCA, the data suggest that DWORF would compete more effectively when calcium is elevated, helping to relieve SERCA inhibition each time that increased transport function is required. In addition, the inverted calcium-dependence of the DWORF-SERCA interaction offers an explanation for the direct activation of SERCA by DWORF, providing a mechanistic framework for the activity data from reconstituted proteoliposomes (**Figure 3**) and the cellular data for SERCA-dependent calcium dynamics (**Figures 4 & 5**). We propose that PLN inhibition involves preferential interaction and stabilization of SERCA enzymatic states that prevail at resting calcium (e.g. (32,41,42)), while DWORF preferentially binds and stabilizes SERCA states that are populated during calcium elevations. The DWORF interaction enhances the kinetics of rate-limiting steps in the SERCA transport cycle.

### Unique structure of DWORF

A critical step was to evaluate the structure of DWORF in comparison to the well-characterized SERCA regulatory subunits PLN (23,32) and SLN (33,34,43). Both SLN and PLN form continuous transmembrane helices with well-defined orientations in the membrane. The cytoplasmic domain of PLN is longer than that found in SLN and DWORF, and it lies along the membrane surface in the structure determined by NMR spectroscopy (PDB code 2KYV (23)). In the X-ray crystal structures of the SERCA-PLN (32) and SERCA-SLN (33,34) complexes, PLN and SLN are found as continuous transmembrane helices (residues ~24-48 and ~1-31, respectively), though the cytoplasmic domain of PLN was not resolved. The helical transmembrane domains of PLN and SLN facilitate the structural interactions in the SERCA-bound states and it is a critical feature of these peptides. In contrast, the DWORF transmembrane domain appears to be discontinuous (residues Leu^17^ to Ser^35^), with a break at Pro^15^ and an N-terminal helix that lies along the membrane surface (residues Met^1^ to Leu^13^).

The molecular structure of the SERCA-DWORF complex remains unknown. The current hypothesis is that DWORF binds to the inhibitory groove and displaces PLN (11), and that DWORF and PLN have similar affinities for SERCA (30). That said, the proposed helix-linker-helix structure of DWORF (**Figure 2**) allows us to speculate about potential mechanisms. The break at Pro^15^ of DWORF occurs at a critical location for PLN and SLN inhibition of SERCA (**Figure 1**). Leu^31^ and Asn^34^ of PLN (Leu^8^ & Asn^11^ in SLN) are two essential residues for SERCA inhibition, and they are positioned to interact with SERCA by the continuous transmembrane helix of PLN. Thus, if DWORF binds to SERCA and replaces PLN, DWORF lacks the N-terminal residues such as Leu^31^ and Asn^34^ of PLN that contribute to SERCA inhibition (**Figure 7A, B**). This model offers an explanation for why DWORF itself does not inhibit SERCA – the discontinuous transmembrane helix and the substitution of a proline residue are inconsistent with structural features known to contribute to SERCA inhibition. The model also offers an explanation for how DWORF activates SERCA. We have previously proposed that modulation of the lipid bilayer by PLN is a mechanism for enhancing SERCA maximal activity (25). DWORF binding to the inhibitory groove of SERCA (**Figure 7B**) would be expected to modulate the lipid bilayer as suggested by the MD simulations (**Figure 2**). Importantly, the thinning of the lipid bilayer observed in the MD simulations is restricted to the cytoplasmic side of the membrane where it could impact the dynamics of transmembrane helices (e.g. M1 & M2) that form the calcium access funnel.

**Figure 7.**
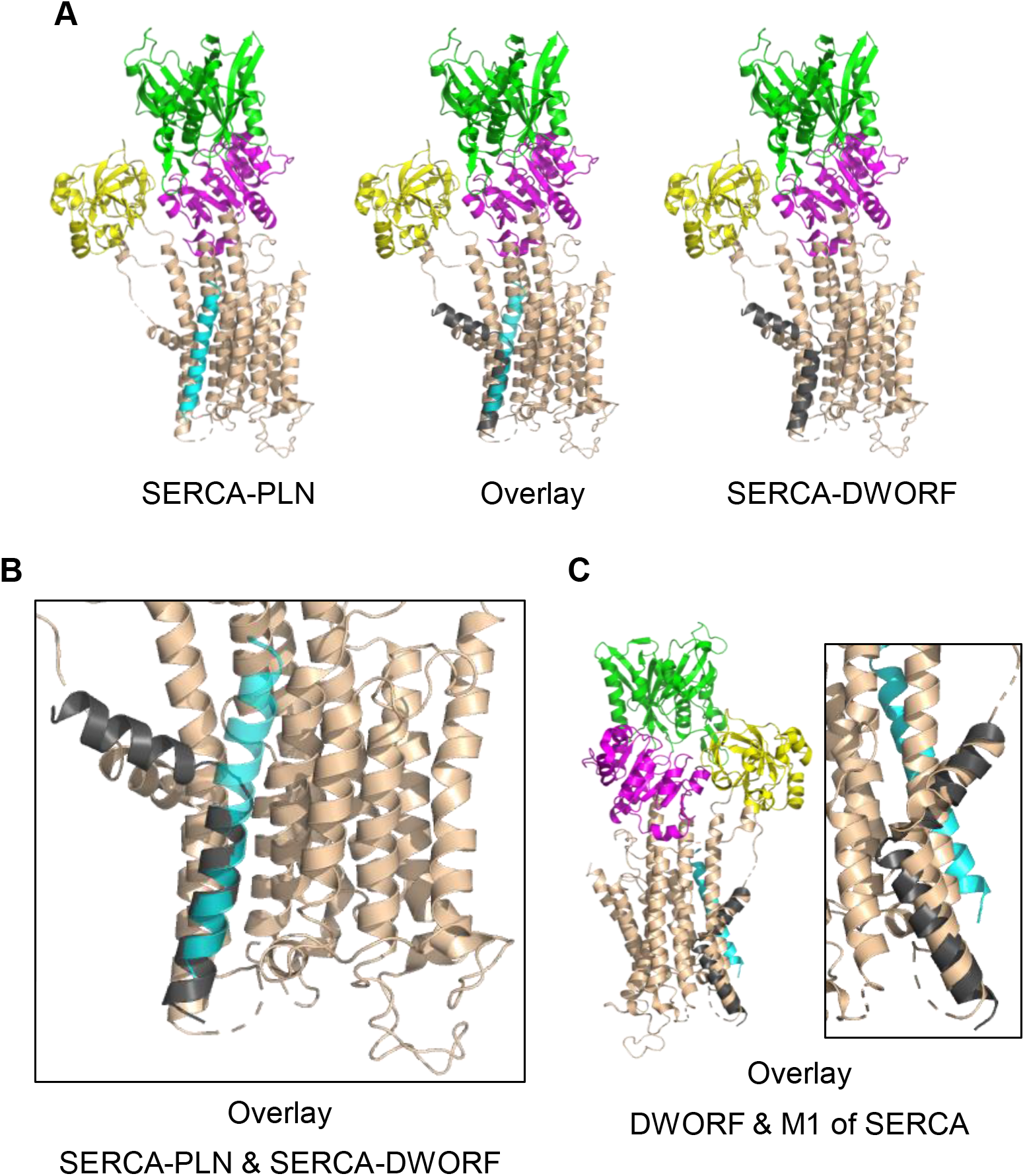
**Figure legend:** Molecular model for the interaction of SERCA with DWORF. (**A**) SERCA-PLN, SERCA-DWORF, and the overlay are shown in cartoon format. The molecular model of DWORF (Figure 2A) was superimposed on the structure of the SERCA-PLN complex (PDB code 4KYT) according to the topological alignment in Figure 1. SERCA is colored tan, with the nucleotide-binding domain in green, the phosphorylation domain in magenta, and the actuator domain in yellow. PLN is shown in cyan and DWORF in grey. (B) Close up view of the SERCA-PLN and SERCA-DWORF complexes. (C) The structure of DWORF in Figure 2A is similar to transmembrane segment M1 of SERCA. Shown is a superimposition of DWORF (grey) and M1 of SERCA (tan).

A surprising outcome of the helix-linker-helix structure of DWORF is that it strongly resembles transmembrane segment M1 of SERCA in the calcium-free conformations (**Figure 7C**). The significance of this is unknown. However, recent structures of SERCA have emerged where transmembrane segments M1 and M2 are more distant from the remaining cluster of transmembrane helices (44,45). By mimicking M1, perhaps DWORF modulates this region of SERCA as a mechanism for enhancing SERCA turnover rate. Finally, another mechanism has been proposed for enhancing SERCA activity, an interaction of the PLN pentamer with M3 of SERCA that increases maximal activity (25). While it is unclear if DWORF is capable of this latter interaction, the data presented herein suggest that DWORF has higher affinity for the calcium-bound conformations of SERCA, which likely alters the E2-E1 transition. The displacement of PLN, the absence of key inhibitory interactions (e.g. Leu^31^ & Asn^34^ of PLN), and perturbation of the membrane bilayer provide potential rationales for the higher maximal activity of SERCA in the presence of DWORF.

## MATERIALS & METHODS

### Molecular modeling of DWORF

MODELLER, a protein structure homology-modeling program (46), was used to generate molecular models of DWORF as a continuous α-helix (residues 1-35) and a helix-linker-helix (residues 1-13 modeled as an α-helix; residues 14-16 as random coil; and residues 17-35 as an α-helix). In evaluating the amino acid sequence of DWORF, we carried out secondary structure prediction using PSIPRED (16), MLRC (17), JPRED v4.0 (18), and Porter v5.0 (19). We carried out transmembrane region prediction using MEMSAT-SVM (16), HMMTOP (20), TMHMM v2.0 (21), and PredictProtein (22). Finally, we carried out hydrophobic moment analysis to identify regions of amphipathic helices using PEPWHEEL (47) and HMOMENT (48).

### Molecular Dynamics Simulations

The DWORF models were inserted in a 1-palmitoyl-2-oleoyl-sn-glycero-3-phosphocholine (POPC) lipid bilayer containing a total 200 lipid molecules using the membrane builder module of CHARMM-GUI web server (49,50). The systems were solvated using a TIP3P water model with a minimum margin of 20 Å between the protein and the edges of the periodic box in the z-axis. Potassium and sodium ions were added to reach a concentration of 150 mM and neutralize the total charge of the system. Molecular dynamics (MD) simulations were carried out using the Amber ff14SB (51) and Lipid 17 force field topologies and parameters implemented in Amber 18 and AmberTools package (52). The systems were energy minimized and equilibrated following the six-step preparation protocol recommended by CHARMM-GUI (53). The temperature was maintained at 310 K with Langevin thermostat algorithm and the pressure was set to 1.0 bar using the Monte Carlo barostat. All bonds involving hydrogens were constrained using the SHAKE algorithm. Each DWORF model was subjected to 2000 ns MD simulation. For the MD analysis, AMBER MD trajectories and coordinates were converted to GROMACS files using MDAnalysis python library (54). We calculated the backbone RMSD of DWORF for the entire trajectory. The frames of each simulation were submitted to a backbone RMSD clustering analysis using a cutoff value of 1.0 Å to retrieve the most representative structure of DWORF.

### Materials

All reagents were of the highest purity available: octaethylene glycol monododecyl ether (C_12_E_8_; Barnet Products, Englewood Cliff, NJ); egg yolk phosphatidylcholine (EYPC), phosphatidylethanolamine (EYPE) and phosphatidic acid (EYPA) (Avanti Polar Lipids, Alabaster, AL); all reagents used in the coupled enzyme assay including NADH, ATP, PEP, lactate dehydrogenase, and pyruvate kinase (Sigma-Aldrich, Oakville, ON Canada).

### Co-reconstitution of DWORF and SERCA

Recombinant human DWORF was expressed as a maltose-binding protein (MBP) fusion with a TEV cleavage site for removal of MBP. DWORF was purified by a combination of organic extraction (chloroform-isopropanol-water) and reverse-phase HPLC (36). Purified DWORF was stored as lyophilized thin films (63 μg aliquots). SERCA1a was purified from rabbit skeletal muscle SR. For co-reconstitution, lyophilized DWORF (63 μg) was suspended in a 100 μl mixture of trifluoroethanol-water (5:1) and mixed with lipids (360 μg EYPC & 40 μg EYPA) from stock chloroform solutions. The peptide-lipid mixture was dried to a thin film under nitrogen gas and desiccated under vacuum overnight. The peptide-lipid mixture was hydrated in buffer (20 mM imidazole pH 7.0; 100 mM NaCl; 0.02% NaN_3_) at 50°C for 15 min, cooled to room temperature, and detergent-solubilized by the addition of C_12_E_8_ (0.2 % final concentration) with vigorous vortexing. Detergent-solubilized SERCA1a was added (300 μg in a total volume of 300 μl) and the reconstitution was stirred gently at room temperature. Detergent was slowly removed by the addition of SM-2 Bio-Beads (Bio-Rad, Hercules, CA) over a 4-hour time course (final weight ratio of 25 Bio-Beads to 1 detergent). Following detergent removal, the reconstitution was centrifuged over a sucrose step-gradient (20% & 50% layers) for 1 h at 100,000*g*. The reconstituted proteoliposomes at the gradient interface were removed, flash-frozen in liquid-nitrogen, and stored at −80 °C. The final molar ratio was 120 lipids to 2 DWORF to 1 SERCA.

### ATPase activity assays of SERCA reconstitutions

ATPase activity of the co-reconstituted proteoliposomes was measured by a coupled-enzyme assay over a range of calcium concentrations from 0.1 μM to 10 μM (37,38). The assay has been adapted to a 96-well format utilizing Synergy 4 (BioTek Instruments) or SpectraMax M3 (Molecular Devices) microplate readers. Data points were collected at 340 nm wavelength, with a well volume of 155 μl containing 10-20 nM SERCA at 30 °C (data points collected every 28-39 seconds for 1 hour). The reactions were initiated by the addition of proteoliposomes to the assay solution. The V_max_ (maximal activity) and K_Ca_ (apparent calcium affinity) were determined based on non-linear least-squares fitting of the activity data to the Hill equation (Sigma Plot software, SPSS Inc., Chicago, IL). Errors were calculated as the standard error of the mean for a minimum of four independent reconstitutions.

### In-cell calcium uptake and spontaneous calcium release methods

pCMV R-CEPIA1er was a gift from Dr. Masamitsu Iino (Addgene, USA). The vector encoding the human RyR2 cDNA fused to GFP at the N-terminus was kindly provided by Dr. Christopher George (University of Cardiff, UK). The vector encoding human SERCA2a cDNA was kindly provided by Dr. David Thomas (University of Minnesota, USA). The SERCA2a cDNA was cloned into the mCerluean-M1 modified plasmid (Addgene, USA), yielding SERCA2a fused to a modified Cerulean fluorescent protein (mCer) at the N-terminus. SERCA2a cDNA was also cloned into the inducible expression vector pcDNA5/FRT/TO for the generation of SERCA2a stable cell line. The sequences were all verified by single pass primer extension analysis (ACGT Inc., USA).

#### Generation of SERCA2a stable cell line

Stable inducible Flp-In T-Rex-293 cell line expressing SERCA2a was generated using the Flp-In T-REx Core Kit (Invitrogen, USA) as described (28). Flp-In T-REx-293 cells were co-transfected with the pOG44 vector encoding the Flp recombinase and the expression vector pcDNA5/FRT/TO containing the SERCA2a cDNA. 48 h after transfection the growth medium was replaced with a selection medium containing 100 μg/ml hygromycin. The hygromycin-resistant cell foci were selected and expanded. Stable cell lines were cultured in high glucose Dulbecco’s modified Eagle’s medium (DMEM) supplemented with 100 units/ml penicillin, 100 mg/ml streptomycin and 10% fetal bovine serum at 5% CO_2_ and 37°C. Expression of SERCA2a in the stable cell line was verified by western blot analysis 48h after induction of recombinant pump expression with 1μg/ml tetracycline. This strategy results in SERCA2a expressed 10-fold over endogenous SERCA2b (28), so calcium uptake is dominated by the exogenous cardiac isoform. MCer-DWORF expression in these cells showed a similar pattern as CEPIA-1er (**Figure 4A**), indicating that DWORF is preferentially localized in the ER membrane. Since CICR depends on RyR2 expression level, we analyzed calcium waves only in cells that had similar GFP-RyR2 signal.

#### Confocal Microscopy

Changes in the luminal ER [Ca^2+^] ([Ca^2+^]_ER_) and cytosolic [Ca^2+^] ([Ca^2+^]_Cyt_) were measured with laser scanning confocal microscopy (Radiance 2000 MP, Bio-Rad, UK) equipped with a ×40 oil-immersion objective lens (N.A.=1.3). T-Rex-293 cells expressing SERCA2a were transiently co-transfected with plasmids containing the cDNA of RyR2, DWORF and R-CEPIA1er. Experiments were conducted 48 h after transfection to obtain the optimal level of recombinant protein expression. The surface membrane was permeabilized with saponin (0.005%). Experiments were conducted after wash out of saponin with a solution of 100 mM K-aspartate, 15 mM KCl, 5 mM KH_2_PO_4_, 5 mM MgATP, 0.35 mM EGTA, 0.22 mM CaCl_2_, 0.75 mM MgCl_2_, 10 mM HEPES (pH 7.2), and 2% dextran (MW: 40,000). Free [Ca^2+^] and [Mg^2+^] of this solution were 200 nM and 1 mM, respectively.

#### [Ca^2+^]_ER_ measurements

[Ca^2+^]_ER_ was recorded as changes in fluorescence intensity of the genetically encoded ER-targeted Ca^2+^ sensor R-CEPIA1er (29). R-CEPIA1er was excited with a 514 nm line of the argon laser and signal was collected at >560 nm. Line scan images were collected at a speed of 10 ms/line. The R-CEPIA1er signal (F) was converted to [Ca^2+^]_ER_ by the following formula: [Ca^2+^]_SE_ = K_d_ × [(F − F_min_)/(F_max_ − F)]. F_max_ was recorded in 5 mM Ca^2+^ and 5 μM ionomycin and F_min_ was recorded after ER Ca^2+^ depletion with 5 mM caffeine. The K_d_ (Ca^2+^ dissociation constant) was 564 μM (3). SERCA-mediated Ca^2+^ uptake was calculated as the first derivative of [Ca^2+^]_ER_ refilling (d[Ca^2+^]_ER_/dt) after RyR2 inhibition with ruthenium red (15 μM) and tetracaine (1 mM). RyR2-independent Ca^2+^ leak was analyzed as the first derivative of [Ca^2+^]_ER_ decline (d[Ca^2+^]_ER_/dt) after simultaneous inhibition of RyR2 and SERCA. ER Ca^2+^ uptake and Ca^2+^ leak rates were plotted as a function of [Ca^2+^]_ER_ to analyze maximum ER Ca^2+^ uptake rate and maximum ER Ca^2+^ load. All 2-D images and line scan measurements for [Ca^2+^]_ER_ were analyzed with ImageJ software (NIH, USA).

#### Cytosolic [Ca^2+^] ([Ca^2+^]_i_) measurements

[Ca^2+^]_i_ was measured in intact cells with the high-affinity Ca^2+^ indicator Fluo-4 (Molecular Probes/Invitrogen, Carlsbad, CA, USA). To load the cytosol with Fluo-4, cells were incubated at room temperature with 10 μM Fluo-4 AM for 15 min in Tyrode solution (140 mM NaCl, 4 mM KCl, 0.5 mM CaCl_2_, 1 mM MgCl_2_, 10 mM glucose, 10 mM HEPES, pH 7.4), followed by a 20 min wash. Fluo-4 was excited with the 488 nm line of an argon laser and the emission signal collected at wavelengths above 515 nm. Spontaneous Ca^2+^ waves were measured at different [Ca^2+^] (0.5, 1, 2, 5 and 10 mM). In the end of each experiment, caffeine was applied to induce maximal ER Ca^2+^ release.

#### Statistics

Data are presented as mean ± standard error of the mean (SEM) of n measurements. Statistical comparisons between groups were performed with the Student’s t test for unpaired data sets. Differences were considered statistically significant at p<0.05. Statistical analysis and graphical representation of averaged data was carried out on OriginPro7.5 software (OriginLab, USA).

### In-cell FRET methods

#### Fluorescence Resonance Energy Transfer Measurements

Acceptor sensitization FRET was quantified as previously described (31). AAV 293 cells were cultured in DMEM cell culture medium supplemented with 10% fetal bovine serum (FBS; ThermoScientific, Waltham, MA). Following culture, cells were transiently transfected using either MBS mammalian transfection kit (Agilent Technologies, Stratagene, La Jolla, CA) or Lipofectamine 3000 transfection kit (Invitrogen, Carlsbad, CA) with either EYFP-PLN or DWORF, and mCerulean (mCer)-SERCA constructs in a 1:5 molar plasmid ratio with the fluorescent protein fused via a 5 amino acid linker to the N-terminus (12,30,55,56). Cells were then plated 24 hours before imaging in 4 well chambered coverglass plates coated with poly-D-lysine and imaged utilizing wide-field fluorescent microscopy. Cells were imaged on a Nikon Eclipse Ti 2 equipped a Photometrix Prime 95B CMOS camera (Tucson, Arizona, USA) and Lumencor Spectra X (Beaverton, Oregon, USA). Imaging was done in a permeabilization buffer containing 120 mM potassium aspartate, 15 mM potassium chloride, 5 mM magnesium ATP, .75 mM magnesium chloride, 2% dextran, 5 mM potassium phosphate, 2 mM EGTA, 20 mM HEPES, and either 0 or 1.7 mM added calcium chloride. Cells were permeabilized in .005% saponin buffer for 1 min and then washed twice with buffer before imaging. Data were collected using a 20x .75 N.A. objective using 50 ms exposure times for all channels and analyzed using in FIJI using a macro selecting cells with a CFP intensity above 150 AU above background, circularity of .4-1 AU, and size of 500-2500 pixels with a rolling background size of 200. FRET efficiency was calculated according to E_app_ = I_DA_ − a(I_AA_)−d(I_DD_) /(I_DA_ − a(I_AA_)+(G−d) I_DD_ (https://www.ncbi.nlm.nih.gov/pmc/articles/PMC1304294/), where I_DA_ is the intensity of fluorescence of acceptor emission with donor excitation; I_AA_ is the intensity of acceptor fluorescence with acceptor excitation; and I_DD_ is the fluorescence intensity of donor emission with donor excitation. G represents the ratio of sensitized emission to the corresponding amount of donor recover during acceptor photobleaching and acts as a correction factor which is constant for a given fluorophore and image conditions. Constants a and d are bleed-through constants calculated from a = I_DA_ / I_AA_ for a control sample transfected with only YFP labeled SERCA, and d = I_DA_ / I_DD_ for a sample transfected with only Cer labeled SERCA. These values were determined to be a = 0.185, d = 0.405, and G = 2.782. FRET intensity of each cell was then compared to the cell’s YFP intensity, which was used as a measure of protein expression([PLN]). This leads to estimates of apparent dissociation constant (K_D_), YFP intensity at ½ FRET_max_, and intrinsic FRET of the SERCA-PLN complex which is equal to the FRET_max_. Data was fit with a hyperbola of the function FRET = (FRET_max_)([PLN])/(K_D_+[PLN]).

## ACKNOWLEDGEMENTS

This research was supported in part through computational resources and services provided by Advanced Research Computing at the University of Michigan, Ann Arbor and Compute Canada (www.computecanada.ca).

## AUTHOR CONTRIBUTIONS

HSY, SLR, and AZ designed the research. MEF, EB, EEC, MPP, and MPD carried out all experiments and analyzed all data. LMEF, RAO, and NR carried out all MD simulations. HSY, MJL, SLR, and AZ interpreted the results and wrote the paper.

